# GRAND: A database of gene regulatory network models across human conditions

**DOI:** 10.1101/2021.06.18.448997

**Authors:** Marouen Ben Guebila, Camila M Lopes-Ramos, Deborah Weighill, Abhijeet Rajendra Sonawane, Rebekka Burkholz, Behrouz Shamsaei, John Platig, Kimberly Glass, Marieke L Kuijjer, John Quackenbush

## Abstract

Gene regulation plays a fundamental role in shaping tissue identity, function, and response to perturbation. Regulatory processes are controlled by complex networks of interacting elements, including transcription factors, miRNAs and their target genes. The structure of these networks helps to determine phenotypes and can ultimately influence the development of disease or response to therapy. We developed GRAND (https://grand.networkmedicine.org) as a database for gene regulatory network models that can be compared between biological states, or used to predict which drugs produce changes in regulatory network structure. The database includes 12,468 genome-scale networks covering 36 human tissues, 28 cancers, 1,378 unperturbed cell lines, as well as 173,013 TF and gene targeting scores for 2,858 small molecule-induced cell line perturbation paired with phenotypic information. GRAND allows the networks to be queried using phenotypic information and visualized using a variety of interactive tools. In addition, it includes a web application that matches disease states to potentially therapeutic small molecule drugs using regulatory network properties.

**Graphical abstract:** Modeling gene regulation across human conditions integrates cancer tissues and cell lines, small molecules, and normal tissue networks.

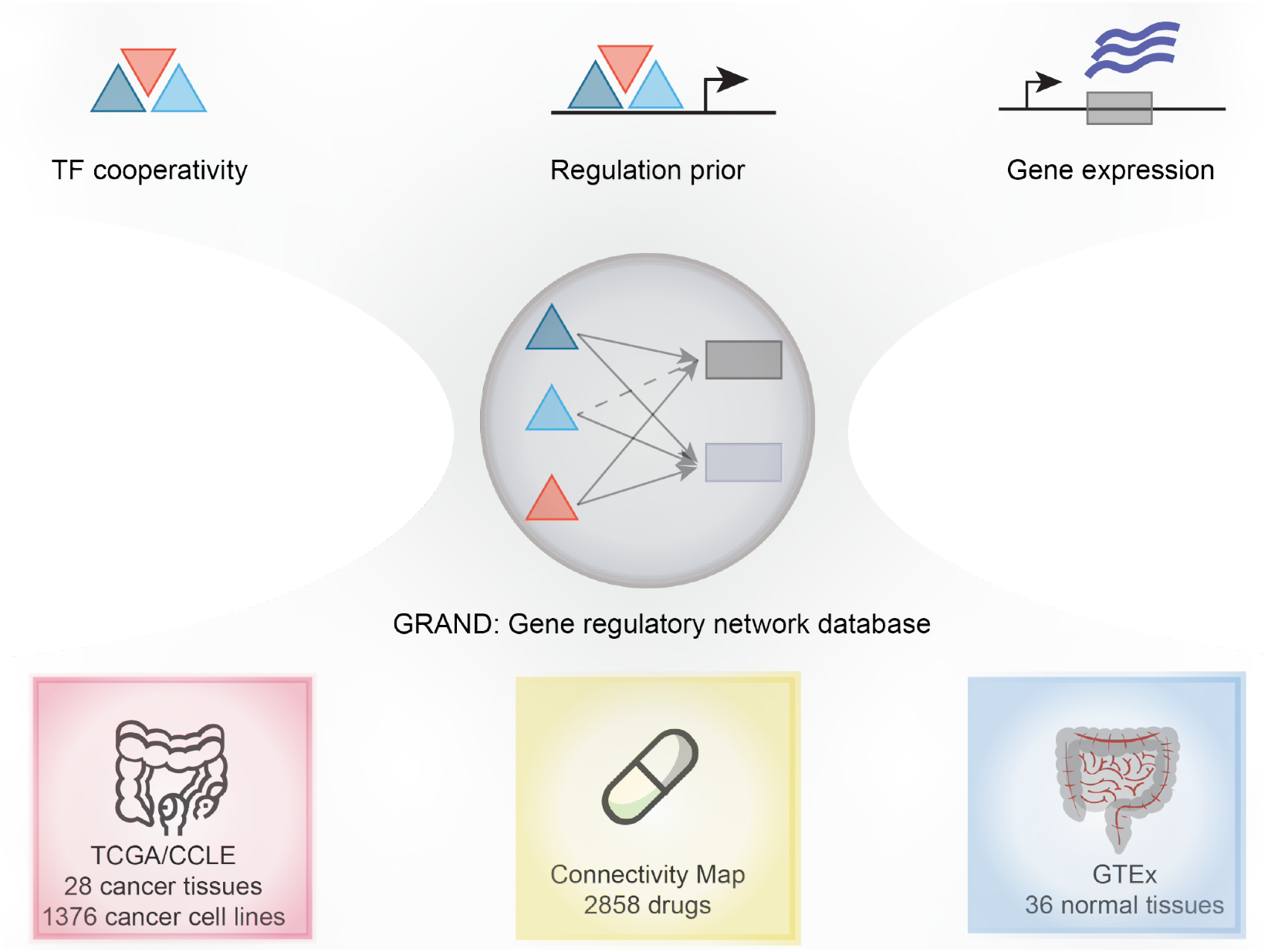

## INTRODUCTION

Gene expression is controlled by complex networks of interacting factors within the cell that help define cellular, tissue, and organismal phenotypes and that allow cells to respond to external and internal perturbations. Dysregulation of these regulatory processes can lead to disease, including cancer (1,2). Although multiple factors play a role in gene regulation (3,4), the most common regulators are transcription factors (TFs) and microRNAs (miRNAs). miRNAs are small noncoding RNAs involved in mRNA post-transcriptional regulation. In most cases, miRNAs bind to short complementary sequences within the 3’ untranslated regions of mRNAs, causing mRNA degradation or translational repression, and thereby silencing their target mRNA (2,5). TFs bind to TF-specific motif sequences in the promoter regions of their target genes and modulate gene expression by interacting or interfering with other key transcriptional proteins including RNA polymerase (4,6). Several experimental techniques such as ChIP-seq (7) and ChEC-seq (8) allow measurement of the binding of TFs across the genome, providing evidence of regulatory associations. However, such experiments typically only look at small numbers of transcription factors and are not scalable to population level studies.

Because large-scale experimental determination of regulatory processes has proven challenging, there is a growing recognition of the need for methods to infer gene regulatory networks (GRNs) and for comparing regulatory network architectures between phenotypes or experimental groups. The rapidly growing volume of genomic and transcriptomic data in human health (9) and disease (10) has greatly facilitated the development of GRN inference methods (11–16) and has provided the validation data necessary to refine and tune these methods. Similarly, the availability of data sets that include both transcriptional profiling and phenotypic response to perturbagens, including small molecule drugs (17–19), provide opportunities to study how expression and regulatory network structures correlate with phenotype. Several GRN databases that provide users with context-specific networks (20–24) have been developed recently. For example, iNetModels (23) has a catalog of coexpression networks in normal and cancer tissues as well as integrated multi-omic networks. GIANT (24) allows to predict tissue-specific networks for a gene of interest and generate hypotheses about functional associations. TRRUST (25) is a curated collection of regulatory interactions that were mined from publications. GRNdb (20) provides a set of regulatory networks using bulk and single-cell data, however, the lack of interactive visualization as well as the lack of availability of the source code of network inference and analysis pipeline could challenge community engagement and reproducibility. The above-mentioned resources were built using approaches that require several gene expression samples to infer context-specific, aggregate GRNs across all samples. However, none of these resources consider sample-specific GRNs to account for essential differences in phenotypic variation between patients such as sex, age, and ethnicity. In particular, none of these resources modeled GRNs in the Cancer Cell Line Encyclopedia (CCLE) database (26), which provides gene expression samples for more than 1,376 cell lines with a single gene expression sample for each cell line. In this case, aggregate methods fail to compute GRNs for individual CCLE cell lines because they require several samples.

Since 2013, our research group has developed and validated a collection of GRN inference tools designed to work with various input data (27–31). This family of tools is collectively referred to as the “Network Zoo” (netzoo; netzoo.github.io). The baseline method in netzoo, PANDA (27), is derived from the understanding that TFs can interact with their target genes to activate or repress the expression of those genes. It also recognizes that some TFs exert their influence as part of multi-TF complexes and that genes that are regulated by the same TFs are likely to exhibit similar patterns of expression. Consequently, PANDA takes as input (i) an initial regulatory network based on mapping TFs to their potential target genes in the genome based on TF binding motifs, as well as (ii) protein-protein interaction (PPI) data, and (iii) the gene co-expression relationships across the samples being studied. PANDA then uses message passing (27) to iteratively search for agreement between these data sources until it arrives at an optimal network structure. This conceptual framework is flexible in that other sources of regulatory information and constraints can be introduced. For example, PUMA (28) extends PANDA by including miRNAs as regulators of expression, while LIONESS (29) uses a linear interpolation approach to extract single-sample networks for each research subject (or biological sample) in a study population. OTTER (30) estimates a gene regulatory network by optimizing graph matching between three networks derived from the three input datasets. DRAGON (31) builds a multi-omic network using a variation of Gaussian Graphical Models (GGMs) by implementing covariance shrinkage to estimate partial correlations.

We previously used the netzoo methods, particularly PANDA and LIONESS, to infer tens of thousands of GRN models. We analyzed these networks in a number of published studies, including GRN comparison of 36 “normal” tissues and two cell lines from the Genotype Tissue Expression (GTEx) project (28,32,33) and six cancers from The Cancer Genome Atlas (TCGA) (30,34–36). Although each study included detailed descriptions of the data and methods used to generate these networks, there was no appropriate data repository for publishing, querying, and visualizing the GRN models themselves due to the large number of genome-scale networks with millions of edges that required more than 6TB of data storage. Given that the inference of these networks took thousands of computational hours, we recognized that the lack of an appropriate network repository to host thousands of network models created substantial obstacles to the reuse of our published network models to investigate additional questions.

To address the need for such a resource and to facilitate the query and analysis of these networks, we created the Gene Regulatory Network Database (GRAND; https://grand.networkmedicine.org). GRAND catalogs curated networks created using netzoo tools together with sample-specific phenotypic information. To supplement the existing collection of networks, and to allow comparison of health and disease phenotypes with perturbations arising from treatment with small molecule candidate therapeutic compounds, we generated additional 173,013 TF and gene targeting scores, which correspond to the weighted outdegree for TFs and weighted indegree for genes (36), from network models of cell lines treated with 2,858 small molecule compounds cataloged by the Connectivity Map (17) project, 1,376 cell line networks from the CCLE database (26) accounting for TF and miRNA regulation, and 22 cancer types from TCGA. In total, GRAND contains 12,468 GRNs representing samples from 36 human tissues, 28 cancer types, 1378 cell lines, and 2,858 small molecule screening assays. The majority of these networks model cis-transcriptional regulation at the TF level, and a subset of networks model post-transcriptional regulation using miRNA information.

The GRNs hosted in GRAND, including the inference pipeline to generate each network, are accessible through an interactive web interface as well as through a well-defined application program interface (API). A network visualization module allows the users to query and plot subnetworks of interest based on several selection parameters as well as to compute the corresponding targeting scores. To support analysis of the collection of networks, we developed two web server applications that allow users to query the GRAND database. The first allows users to perform functional enrichment analysis on a set of TFs ranked by targeting score. The second utility is similar to Connectivity Map analysis (17), but uses network features instead of expression to identify candidate drugs and drug combinations that could be used to reverse or alter regulatory patterns in a particular disease state. Finally, users can upload their own networks for visualization and analysis in GRAND. To demonstrate the utility of GRAND, we present an example in which we compare GRNs between colon cancer and normal colon tissue to identify the genes that are differentially targeted by key regulatory TFs. We then use these to identify an investigational drug that may have a specific effect in colon cancer.

GRAND is a large-scale, multi-study catalog of GRNs that provides regulatory models for perturbed and unperturbed human cell lines, as well as normal and cancer tissues. Our goal is to continue to grow both the number and diversity of network types in GRAND as the field of GRN inference evolves and to add new analytical tools as more phenotypes and experimental samples become publicly available.

## DATA COLLECTION AND DATABASE CONTENT

### Overview of network models in GRAND

GRNs in GRAND are built on the conceptual framework first presented in PANDA in which we model GRNs explicitly as the interaction between TFs and their target genes (Figure 1A). GRAND includes additional network inference tools to model the regulation between miRNAs and their target genes (PUMA), to build single-sample GRNs (LIONESS), to construct GRNs using relaxed graph matching (OTTER), and to use Gaussian Graphical Models to build multi-omic networks (DRAGON). Our starting point in assembling GRAND was the collection of network models we had previously constructed using data from GTEx, TCGA, and GEO (28,30,32,33,37) (Figure 1B-C). To these, we added network models inferred using data available from the Connectivity Map (CMAP) project (17) and CCLE (26). The CMAP project measured gene expression in human cell lines after exposure to a combination of 2,858 approved and investigational drugs and additional chemical compounds. The CCLE collected multi-omic data—miRNA and gene expression, methylation, histone marks, and protein levels—for more than 1000 cell lines (Table S1). These networks can be selected using phenotypic information (Figure 2) and visualized on the browser using a dedicated module (Figure 3).

**Figure 1.**
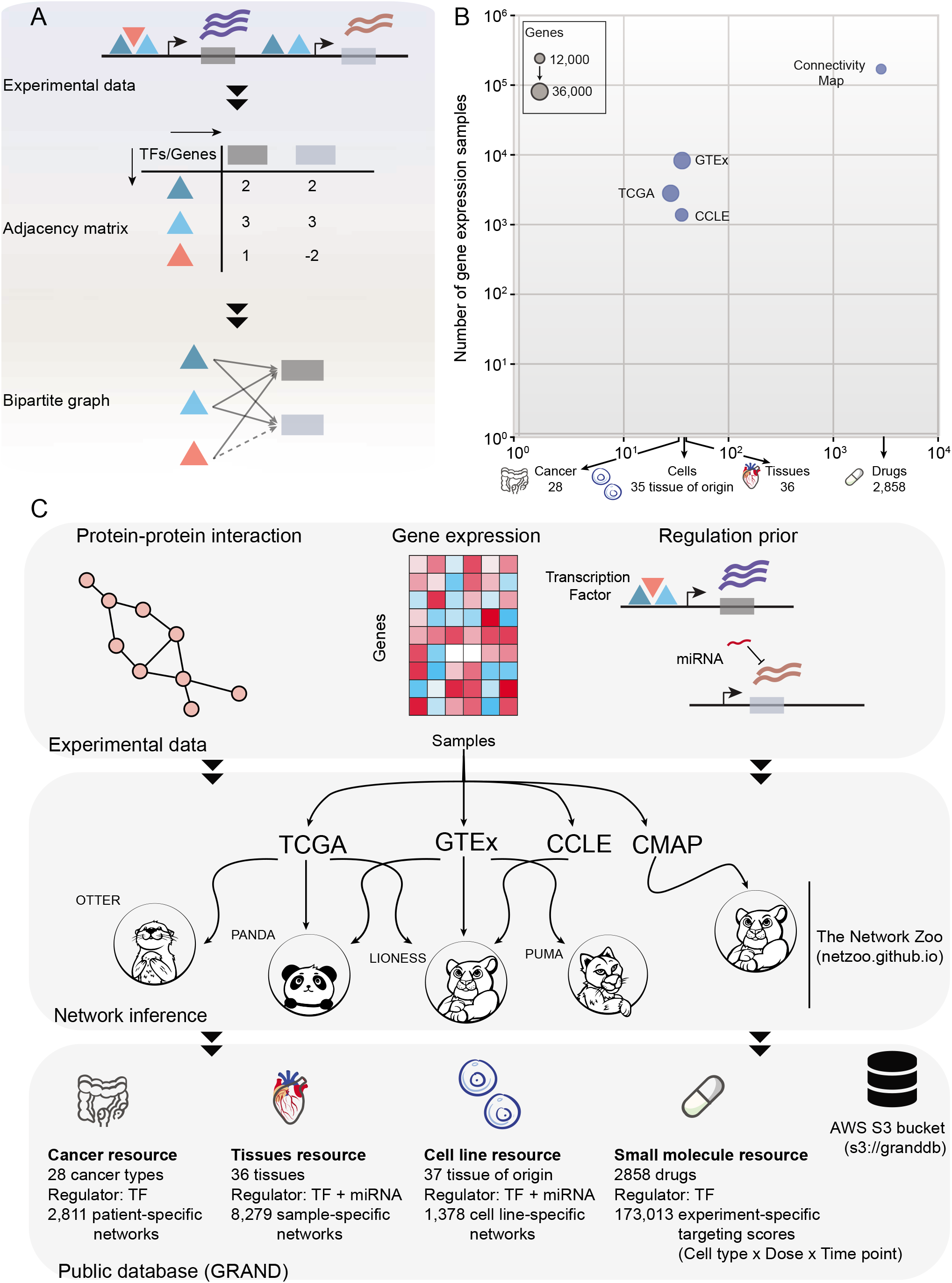
GRAND database statistics and network reconstruction pipeline. **A.** Regulators (TFs) bind in the promoter region of target genes and affect their expression, which can be represented as a bipartite graph and its adjacency matrix. **B.** Representation of the largest gene expression datasets in each of the GRAND resources. X-axis indicates the number of cancer types, tissues types, cell line tissues of origin, and drugs in each dataset. Y-axis indicates the number of samples used to build the networks. The bubble size is scaled by the number of genes in the networks. **C.** GRNs were inferred from experimental data priors such as protein-protein interaction, gene expression, and regulatory prior build from TF motifs or miRNAs predicted targets. The network inference methods that were used are available at netzoo.github.io.

**Figure 2.**
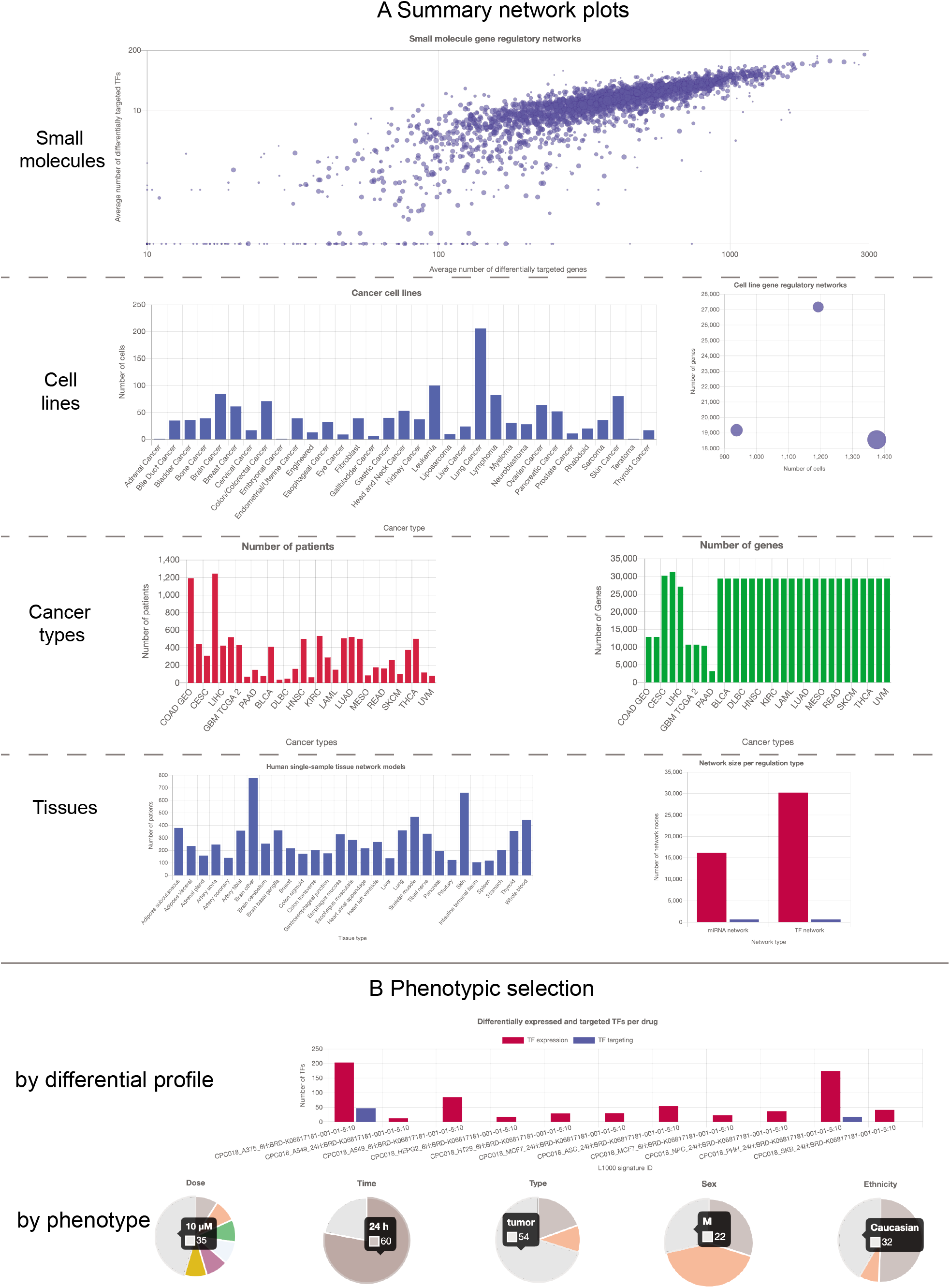
Screenshot of small molecule, cell line, cancer types, and tissues summary plots in GRAND. **A.** The main page for each resource displays a summary interactive plot for the catalog of networks. For small molecules, a bubble plot for each compound leads to the targeting scores across doses, cell lines and sampling times. Cell line, tissue, and cancer TF and miRNA networks are organized by tissue of origin. **B.** A sample-specific network can be selected interactively by differential expression or targeting score of TFs and genes or by phenotypic variables such as donor age, sex, and ethnicity.

**Figure 3.**
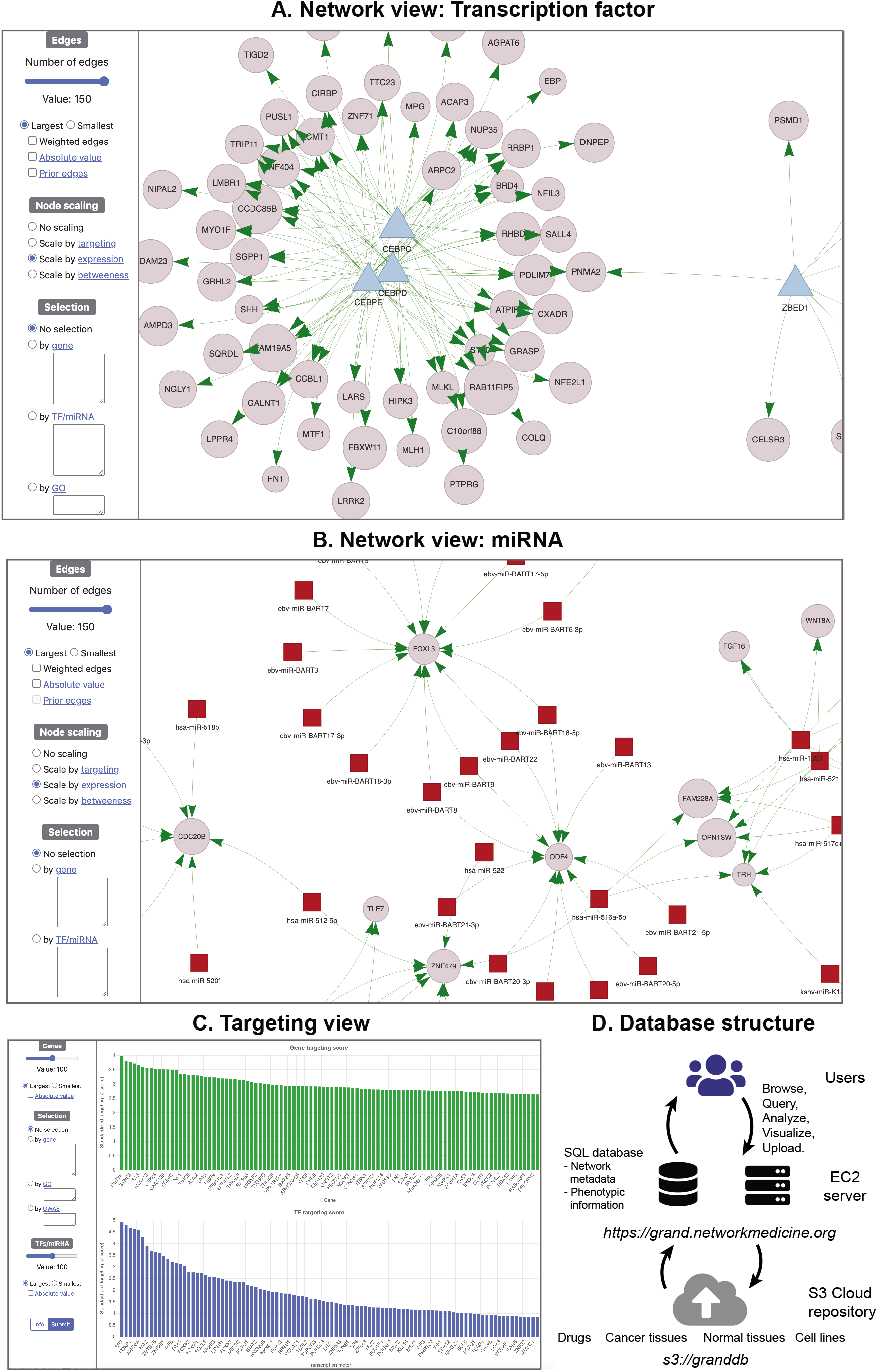
Gene regulatory network visualization and analysis in GRAND. Any network in GRAND can be visualized; shown in this figure are a TF GRN **(A)** and a miRNA GRN **(B)**. Users can select a subset of the network using several parameters related to the edges or the nodes, such as regulators and gene sets, GO terms, and GWAS traits. Nodes can be scaled by expression, targeting or betweenness. **C.** The targeting analysis allows users to calculate and visualize each network’s TF and gene targeting score, and contains links to GRAND’s downstream analysis tools such as functional enrichment analysis and drug repurposing. **D.** Database design and infrastructure.

## GENE REGULATORY NETWORKS

### Small molecule resource

The Connectivity Map phase I (17) and phase II (19) amassed gene expression profiles for human cell lines exposed to various drugs and drug candidates; we selected 2,858 that were cataloged in the Drug Repurposing Hub (DRH) (38). The DRH has essential information on compounds that includes drug indication, chemical structure, and targets. This provided 173,013 gene expression profiles (level 4) for drug exposure across normal and cancer cell lines, doses, and sampling times that were used for GRNs reconstruction (Figure S1).

The Connectivity Map directly profiles the expression of 1,000 genes (the L1000 genes) and uses these data to infer the expression levels of the remaining genes. For network inference, we used the complete set of 12,328 sequenced and inferred genes (https://grand.networkmedicine.org/genes/), also referred to as All Inferred Genes (AIG) set. For these data, we used GPU-accelerated MATLAB implementations of PANDA and LIONESS in the netzoo package (netZooM v 0.5.1) to infer sample-specific GRNs for each of the 173,013 profiles, and subsequently computed TF and gene targeting scores for each network.

### Cancer resource

The cancer resource in GRAND includes both aggregate networks and patient-specific networks across 28 cancer types. In total, 2,811 patient-specific networks were generated for colon cancer, pancreatic cancer, and glioblastoma. The colon GRNs were derived using expression data for 12,817 genes from 445 samples in TCGA and 1,193 samples found in GEO as described previously (34) (Figure S2). Glioblastoma networks were generated on 10,701 genes from 953 samples in TCGA and 70 samples across 10,439 genes from the German Glioma Network (GGN) (35). Pancreatic cancer networks were generated from 150 samples from TCGA spanning both basal-like and classical subtypes, across 3,214 genes (36).

We used PANDA to generate aggregate networks for 22 cancer types in TCGA, and OTTER to generate networks for 3 cancer types in TCGA (30) that were used to validate the accuracy of this new inference tool (30). OTTER is built on the same conceptual framework as PANDA but formalizes the network inference as an optimization problem that maximizes the matching between the three prior graphs representing the input TF-gene regulatory network, TF-TF PPI networks, and correlation-based networks derived using gene expression data. OTTER breast cancer networks include 31,247 genes and represent 1,134 tumor samples. The cervical cancer networks include 30,181 genes and represent 306 tumor samples. The liver cancer networks include 27,081 genes from 374 tumor samples. The validation of these specific networks using ChIP-seq data from ReMap (7) as described by Weighill et al. (30) was added in the “Network Benchmarking” section. In addition, we added the benchmarks of PANDA and DRAGON in the “help” page.

### Tissue resource

The tissue resource made use of GTEx data to construct TF and miRNA GRNs for 36 “normal” human tissues (Figure S2). We used PANDA to build the aggregate TF networks (33), and PUMA to build the aggregate miRNA networks (28). Using PANDA and LIONESS, we also built 8,279 sample-specific TF networks (37).

### Cell line resource

The cell line resource includes TF and miRNA aggregate networks built using PANDA (32) and PUMA (28), respectively, for LCLs and fibroblasts in the GTEx data. Using DRAGON, we also generated an aggregate miRNA network from the 938 CCLE cell lines that had both miRNA and gene expression measurements. Finally, we generated 1,376 single-sample TF networks with LIONESS using CCLE gene expression data from the 1,376 cell lines that had gene expression data corresponding to 35 cancer types.

## ANALYSIS TOOLS IN GRAND

### Finding small molecule candidates through reverse gene targeting

The hypothesis underlying our GRN analysis is that changes in the targeting of genes by TFs represents regulatory differences that underlie phenotypic diversity, including the potential to respond to particular stimuli. These analyses generally search for differentially targeted genes or differential targeting by TFs and use functional enrichment analysis to explore functional differences between the biological states that are compared. In GRAND, we implemented a method, CLUEreg, to extend this framework to the identification of drugs that can potentially reverse disease phenotypes by allowing users to search for regulatory changes induced by small molecule compounds and other drugs profiled in the Connectivity Map.

To build CLUEreg, we extended the small molecule resource in GRAND to all the approved and experimental drugs profiled in the Connectivity Map, consisting of 19,791 total small molecules. Because each small molecule is administered to multiple cell lines using a variety of doses and sampling times, we used PANDA to build an aggregate GRN for each drug, totaling 19,791 GRNs. For each drug-specific GRN, we constructed a “targeting score” for each gene as the sum of inbound edge weights. For each TF, we calculated a targeting score as the sum of outbound edge weights. The targeting score of all drug-specific GRNs are assembled into a gene-by-drug or TF-by-drug targeting matrix. We then reduce the complexity of these matrices to the set of “differentially targeted/targeting” genes or TFs by comparing the targeting weight to the distribution of weights within the matrix and selecting as differentially targeted/targeting those genes/TFs that have targeting scores that deviate with more than two standard deviations from the mean.

To use CLUEreg, users provide two lists, one consisting of genes (or TFs) with increased targeting and the second consisting of genes (or TFs) with decreased targeting in the disease of interest. These are compared to the library of 19,791 drug-specific GRNs to identify small molecule drug treatments that likely reverse the targeting score of the gene/TF in the original input GRN. For a given input of a differentially targeted gene (or TF) list, CLUEreg computes two measures of agreement with the effect of each drug (Figure 4).

**Figure 4.**
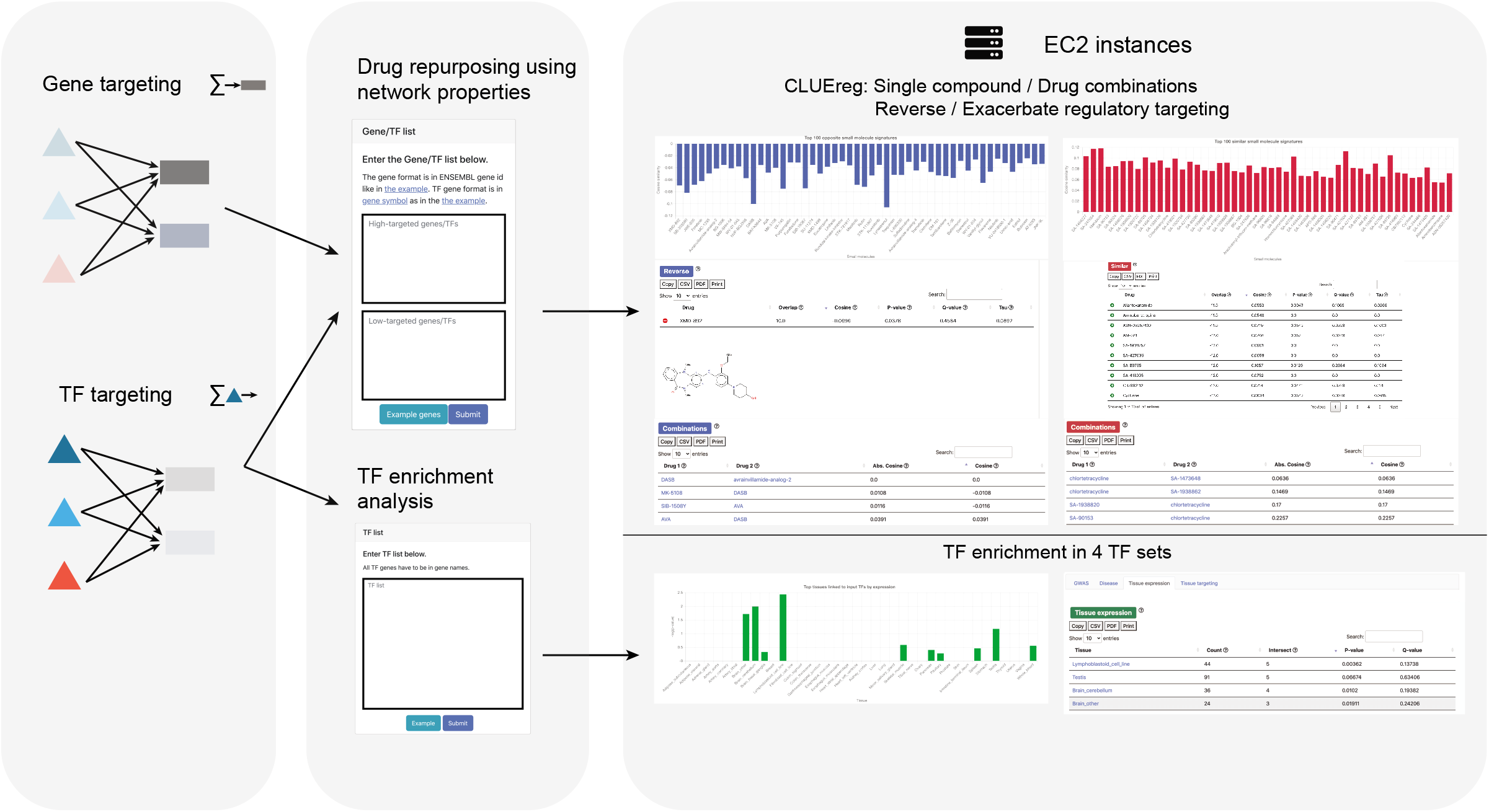
Analysis tools and the web server functionalities in GRAND. A list of up-targeted and down-targeted genes or TFs computed from a weighted bipartite network are given as an input to CLUEreg, which then computes similarity scores to the targeting scores of 19,791 small molecules to find the single and combination candidates that reverse or exacerbate the input signature. A second feature allows users to perform an enrichment analysis of a list of TFs against four TF sets: TFs linked to disease phenotypes through GWAS or the Human Phenotype Ontology, and differentially expressed or differentially targeting TFs in specific tissues.

The first is the cosine similarity comparing the differentially targeted gene lists in a user’s input query and a specific drug as described in Duan et al. (39) and defined as:

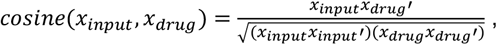

Where *x_input_* denotes the user input vector of differentially targeted genes (or TFs) and *x_drug_* the vector of differentially targeted genes (or TFs) for a given drug. A cosine similarity equal to −1 indicates that the drug has a regulatory pattern that is the reverse of the input list, suggesting that the drug is a candidate for reversing the differential regulation induced by the disease state under investigation. In contrast, a cosine similarity equal to 1, indicates that the small molecule exacerbates the input list, as it aligns perfectly with its direction and sense.

The second measure computed by CLUEreg is the overlap score (39) between the input list and the differentially targeted genes (or TFs) for each drug, defined as:

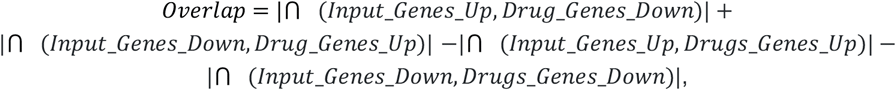

A positive overlap score between a query targeting list and a drug targeting list suggests that the drug reverses the input, while a negative overlap score suggests that the drug and the input have similar regulatory effects. In developing the application, we have found that both metrics provide highly consistent rankings of candidate drugs.

CLUEreg computes a p-value for each drug candidate by resampling 10,000 random inputs of varying lengths as a null distribution. In addition, q-values are provided as corrected p-values using the Benjamini-Hochberg procedure (40). There are several drug classes that can induce profound changes on transcription and often produce false positives in connectivity analysis. An example of such drugs are Histone Deacetylase (HDAC) inhibitors. To control for these effects, we computed a tau-value as described in the Connectivity Map (17). First, we computed the cosine similarity of each drug in CLUEreg against all other drugs to generate a cosine distribution. Then to generate the tau-value, we rank the cosine between the input query and a given drug within the precomputed distribution of all drugs. Tau varies between 0 and 1 and represents the fraction of drugs in the database that have a stronger connectivity. Low tau-values indicate specific activities, while large tau values indicate compounds with promiscuous effects.

We also implemented drug combinations in CLUEreg as described in Duan et al. (39) by ranking pairs of drugs within the top 20 hits. Drug pairs are ranked by their cosine similarity such that an optimal pair has a cosine of 0, which indicates activity on orthogonal gene/TF vectors. Therefore, the optimal drug combination has compounds that optimally reverse the input regulatory profiles while acting on different target genes and pathways.

### TF enrichment analysis tool

Comparative gene regulatory network analysis generally identifies “differential targeting” TFs that regulate different sets of genes in the phenotypes being compared. To help characterize sets of TFs, GRAND implements a hypergeometric test to compare a user-supplied list of TFs to a variety of resources, including a list of tissue-specific differential targeting and differentially expressed TFs (33), a library of 170 GWAS traits in which a GWAS SNP maps to a TF’s corresponding gene (6), and a collection of TFs identified by the Human Phenotype Ontology (41) library that includes 2,440 human conditions and phenotypes. The tool computes the p-value and the multiple testing corrected q-value to assess the significance of the enrichment of the term in the input TF query in the background of 1,639 TFs encoded in the genome (Figure 4).

## DATABASE CONSTRUCTION AND USER INTERFACE

### Database structure, design, and implementation

The GRAND frontend was developed in Bootstrap (v 5.0) and jQuery (v 3.3.1). Network visualization was implemented in Vis.js (v 8.5.2). Bar plots, scatter charts, and bubble plots were implemented using Chart.js (v 2.9.4) and Highcharts.js (v 8.2.2). The backend was developed in Django (v 3.0.5) (42) and Python (v 3.8) (43) and deployed on a Ubuntu (v 18.04) Amazon Web Services (AWS) EC2 instance using Nginx (44) web server and SQLite (v 3.31.1) database tool which is integrated in Django (Figure 3D). Using Django for constructing the website was motivated by its versatility as it integrates a frontend tool, a database management system, and a backend tool, which provides great ease-of-use.

GRAND contains more than 6 TB of network data which is hosted on a public AWS S3 bucket (s3://granddb). Although websites such as NDEx (21) allow users to host and visualize networks for up to 10 GB of data, the size and complexity of data in GRAND required a tailored design approach to efficiently process queries on genome-scale networks with millions of edges. Finally, programmatic access to the website through the API was implemented using Django REST Framework (v 3.11). The website repository is version-controlled at https://github.com/QuackenbushLab/grand.

### User interface: Network Browsing

GRAND’s interface was designed to allow users to browse, download, visualize, and analyze the collected set of networks. The networks are organized by source type and include links from the homepage to browsable sets of network models from “Small molecules,” “Cancer,” “Tissues,” and “Cell lines”; these pages can also be reached using the “Networks” menu item in the upper right menu bar. Each page contains multiple links to brief “help” messages that explain various fields. Clicking on one of these collections takes the user to a subpage where the subsets of the main classes can be selected. Drug targeting scores are classified by the drug name, with an interactive bubble plot that provides information about the differentially targeted TFs and genes as well as the number of samples in each drug. The “Cancer” page classifies cancer types by tissue of origin. Three bar plots summarize the number of samples, TFs, and genes in each network and allow users to access the cancer type of interest by clicking on bars within the plots (Figure 2A). The “Tissues” page lists all 36 tissues in a data table. A bar plot summarizes the number of sample-specific networks available in each category (Figure 2A). A second bar plot categorizes networks by regulation modality (TF or miRNA). These plots are interactive and clicking on individual bars filters the table below. The “Cell lines” page contains networks categorized into three sets: cancer cell line networks from CCLE, normal cell line networks from GTEx, and a miRNA aggregate network. Cancer cell lines are grouped by cancer type and an interactive bar plot lists the number of samples in each category (Figure 2A). A second, interactive bubble plot shows the size (number of TFs, miRNA, genes, and sample) in each of the three sets.

Clicking on a cell line/cancer/tissue link within these summary pages leads to an individual network page that lists available networks for the given category. In addition, the page provides sortable metadata used for network inference as well as additional metadata, including basic statistics on the type and number of regulators, genes, and samples used to reconstruct the network. In the “Cancer” and “Tissues” sections, the sample number links to the phenotypic variables associated with each sample (Figure 2B). In the “Cell line” and “Small molecules” sections, information is provided on the cell line and drug dosage as appropriate. In the “Small molecules” page, clicking the “Genes” column opens a table containing the gene names and their attributes. In all pages, clicking on the entry in the “Reference” column either links to the relevant published study, or, for the “Small molecules” page, to the relevant entry in PubChem. Each drug in the “Small molecules” section includes a panel with information about the drug indication, its chemical structure, and several relevant parameters compiled from the DRH (38) and the Connectivity Map (17) (Figure S1).

In addition to the network information page, relevant metadata about the samples used in the analysis are available in the “Phenotypic information” table. For aggregate networks, this table was intended to give information about the samples used for network reconstruction and classify the samples by variables such as sex, age, ethnicity, and survival. For single-sample networks, the phenotypic information page allows the user to visualize and download the sample-specific network. To facilitate the selection of networks, phenotypic variables are classified into continuous variables, such as height and age, and categorical variables, such as sex and ethnicity.

Continuous variables are plotted as scatter plots at the top of the network page. Clicking on an individual sample within the plot links to the network visualization page. When continuous variables are missing, we display additional information about the data such as the number of differentially targeted and expressed genes and TFs in each cell line and drug sample, as well as the top ten enriched GO terms for differential TF and gene targeting in cancer. Categorical variables are plotted using pie charts in the phenotypic information page. Clicking on each category within individual pie charts filters the phenotypic information page by the selected phenotype.

The networks and associated metadata can be downloaded, either in bulk or individually, from both the web interface and the API. Users can specify whether to download the networks as either TF-by-gene adjacency matrices using the “Adj” button or lists of TF-gene edges using the “Edge” button. The “Vis” button links to the integrated visualization module that allows users to produce interactive graphs of regulatory networks (see the section on network visualization below).

Finally, reflecting our commitment to reproducible research, clicking on the “Code” button in each network links to the code used to generate the networks along with information about the parameters used in the analysis. For networks generated using MATLAB, the code is provided as “.m” files, while for Python and R, Jupyter notebooks are provided that can be run through the webserver “netbooks” (http://netbooks.networkmedicine.org).

### User interface: Network Visualization

The network visualization tool can be accessed through the “Vis” button in the network table and through the phenotypic variable plots. The network visualization page contains a “network” tab and a “targeting” tab. The “network” tab has a selection panel that allows users to plot a TF (Figure 3A) or miRNA (Figure 3B) subnetwork using several parameters, such as the number of edges and edge weights filtered by absolute or signed values. The “Prior” edges option plots network edges supported by the presence of a TF motif in the promoter region of target genes or miRNA target predictions. Node sizes can be scaled by the targeting score of each node, the average gene expression of the node, or the betweenness centrality of each node in the subnetwork. A regulator (TF or miRNA) and gene list submission form allows users to enter a gene or TF list of interest in both ENSEMBL gene ids and gene symbols to be selected in the network view. An additional GWAS form allows selection of genes by GWAS traits from the GWAS catalog (45). A GO term form allows input of GO terms to select a subnetwork of the term of interest.

Clicking on the “submit” button retrieves the network from the cloud repository and plots the corresponding graphs within the browser interface. Once plotted, the network can be further dynamically manipulated using several options in the configuration panel to change the layout and colors. The network plot can be exported as a file using the “save” button. A network table containing the source and target node names and the edge weight can be downloaded in the bottom panel and a network information section provides basic information about the network with a button that redirects to the full network information page. The “targeting” tab (Figure 3C) computes gene and TF targeting scores in the network and allows selection based on the same parameters as in the network tab. In addition, after plotting targeting scores for the nodes of interest, an analysis section redirects the user to downstream analysis tools such as CLUEreg, for drug repurposing, or TF enrichment analysis, with prefilled forms.

### User interface: Network Analysis

The “Analysis” section provides access to four web server tools: CLUEreg, TF enrichment analysis, network comparison, and visualization and integrated analyses of user-provided networks (Figure 4). While CLUE (CMap and LINCS Unified Environment; https://clue.io) (17) uses gene expression to match drug perturbations to input disease gene lists, CLUEreg uses the properties of inferred regulatory networks to identify drugs that may “correct” aberrant regulatory patterns. The CLUEreg page provides two panels allowing users to enter lists of “high-targeted” and “low-targeted” genes or TFs in the disease of interest. Users can query by gene symbols, ENSEMBL gene ids or mixed lists, by target genes or TFs, and by including or excluding investigatory drugs. An additional option computes optimal drug combinations. CLUEreg outputs the top small molecules that either reverse or enhance the differential targeting in disease, including summary statistics (cosine, overlap, p-value, q-value, and tau-value described in the “Finding small molecule candidates through reverse gene targeting” section). Each row in the result table has an “expand” button that shows the chemical structure and basic information about the drug. The results are also displayed as an interactive bar plot. Clicking on the plot filters the result table for the compound of interest.

The TF enrichment analysis allows users to input a set of TFs in gene symbol, ENSEMBL gene ids or mixed lists and test the enrichment against four TF sets: TFs linked to disease phenotypes through GWAS (6), TFs annotated to disease through the Human Phenotype Ontology (41), and TFs that have previously been identified as either differentially expressed or differentially targeting in specific tissues (33). The results are presented in interactive bar plots and tables showing the enrichment statistics (p-value and q-values).

The “Upload your own network” tab allows users to upload an adjacency matrix as a file of 500 Mb maximum and visualizes the network using an integrated module, perform differential targeting analyses, and export the results to either CLUEreg or Enrichment analysis using prefilled forms.

In addition to using CLUEreg and TF enrichment tools on user-provided gene lists, these tools can be used on any network in GRAND. From the visualization page of a given network, users can run these downstream analyses on a subnetwork of interest. Finally, in the “Network comparison” tab, differential network analyses can be performed on a set of cancer and normal tissues to find regulatory disruptions involved in malignant processes. These networks were generated using the same gene expression and network inference pipeline to remove variability due to parameter choice.

### Additional information and API

GRAND includes a “Help” page that contains extensive information detailing the various sections of the website including a “Contact” section allowing questions to be directed to the website administrators. The help page contains summary information about the data sources and the GRN inference tools. The benchmark section includes bar plots of the benchmarking results of PANDA and DRAGON as they were described in their original publications. Cancer-specific OTTER benchmarks were added in the breast, liver, and cervix cancer pages. Finally, the GitHub link redirects users to the repository containing the code for the website.

Programmatic access to GRAND networks is enabled through an API implemented using Django REST Framework to allow batch downloads and integration into computational pipelines. In addition, the server-side filtering functionality allows users to programmatically select the networks based on a set of query parameters. The API functions and documentation, as well as two tutorials in Python and MATLAB are provided in https://grand.networkmedicine.org/help/#api.

## EXAMPLE ANALYSIS: COMPARING COLON CANCER AND NORMAL COLON NETWORKS

To demonstrate the use of GRAND, we compared networks from modeled colon cancer and normal colon tissues to identify differentially targeted genes in cancer and to suggest small molecules that can potentially reverse the disease-specific network targeting scores. We compared an aggregate PANDA network for colon cancer (34) and the corresponding normal tissue network (33) that had been published using data from TCGA (10) and GTEx (9), respectively. We pruned each network to include only the 12,817 genes and 661 TFs appearing in both.

To compare these networks, we simply subtracted the cancer network from the normal network (Figure 5A). We calculated a targeting score for the genes and TFs as the sum of the weighted in-degree or out-degree, respectively. The genes and TFs were ranked by their respective weights. The 300 genes with the highest and 300 genes with the lowest weights in the differential network were selected for analysis in GRAND; similarly, the 100 highest and the 100 lowest targeting TFs were selected (Figure 5B). We analyzed these gene and TF sets using CLUEreg.

**Figure 5.**
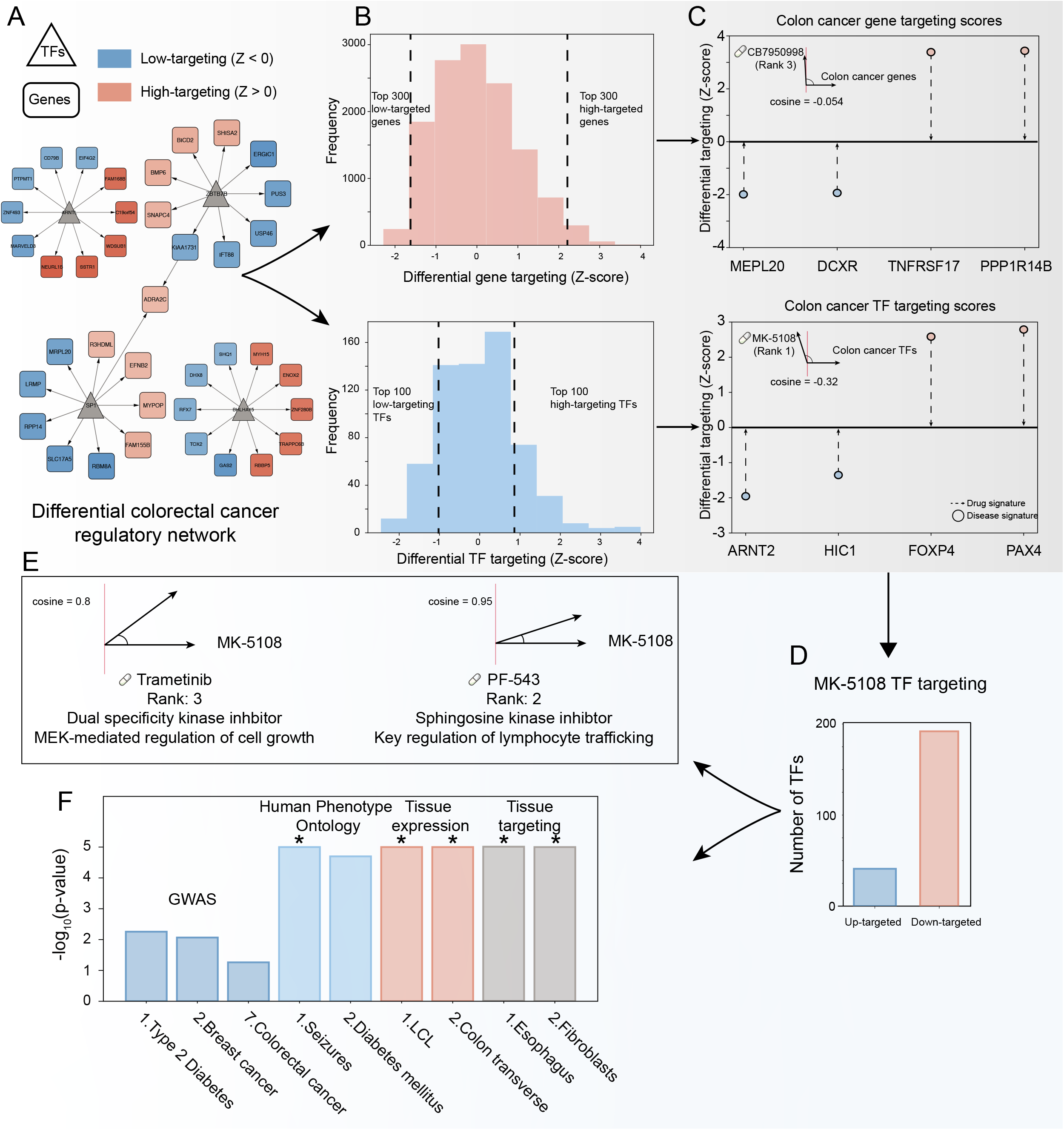
Integrative analysis of colon cancer network using GRAND combined tools. **A.** A differential network between the colon cancer network and the normal transverse colon network allows the selection of the top differential targeted genes and the top differential targeting TFs **(B)**. **C.** CLUEreg analysis suggested two compounds MK-5108 and CB7950998 to reverse the colon cancer network targeting score. **D.** The TF targeting scores of MK-5108, an investigational kinase inhibitor, is similar to the scores of two other known kinase inhibitors **(E)**. Both kinases have different physiological roles which could set the basis for a combination therapy. **F.** TF enrichment analysis of MK-5108 TF targeting scores suggested a possible specificity for colon tissue. * p-value < 10^-5^.

CLUEreg identified a number of drugs as candidates likely to reverse the differentially targeted genes scores in colon cancer. The known anti-cancer compound CB7950998 was among the highest-ranked (rank 3 overall with a cosine of −0.054); in particular CB7950998 was predicted to reverse the targeting of *DCXR* and *MPL20* (Figure 5C), two genes known to be dysregulated in colon cancer. CB7950998 has been suggested to increase the chemosensitivity through acting as *AHR* agonist, however with limited activity *in vivo* (46).

In analyzing the TF targeting scores, CLUEreg identified MK-5108 (rank 1 with a cosine of −0.32) (Figure 5C) as the most likely drug to reverse regulatory targeting in colon cancer and suggests that it works primarily by targeting transcription factor *FOXP4*. MK-5108 is an investigational drug that targets aurora A kinase, a proliferation marker (47) that plays a central role in mitosis (48). Using GRAND to search for the regulatory pattern of MK-5108, we find that the drug is associated with 192 low-targeting TFs and 41 high-targeting TFs (Figure 5D). We then used these TFs as input to CLUEreg to search for compounds with similar targeting patterns. This identified PF-543, a sphingosine kinase inhibitor that alters lymphocyte trafficking (Figure 5E) (49), and Trametinib, an inhibitor of MEK1 and MEK2 that has shown promise in clinical trials for colorectal cancer (50) and metastatic melanoma (51) carrying the BRAF V600E mutation (51).

To further investigate the potential activity of MK-5108, we analyzed the functional roles of the TFs using the TF enrichment tool in GRAND. Searching the list of 233 TFs against the GWAS hits library, type 2 diabetes, breast cancer, and colorectal cancer were identified as the first, second, and seventh most significant GWAS traits (Figure 5F). The search against the Human Phenotype Ontology identified diabetes and seizures as the top traits associated with MK-5108, which may indicate that these could be possible adverse reactions associated with MK-5108. The search of the MK-5108 against the “normal” tissue expression and tissue targeting identified an association with transverse colon tissue as well as the lymphoblast and fibroblast cell lines. The former is logical as MK-5108 is predicted to be effective against colon cancer, the latter cell lines also make sense because MK-5108 targets the mitotic process and these cell lines are known to have altered cell cycle processes relative to their tissues of origin.

While only suggestive and requiring validation experiments, the lines of evidence from multiple sources suggest that MK-5108 may be an agent with efficacy in treating colon cancer by altering regulatory patterns in the disease. More importantly, this example demonstrates the potential value of the GRAND database and its associated search tools and underscores the value of methods for gene regulatory network inference.

### Conclusions and future development

An increasing number of studies involves the inference of GRNs and their subsequent analysis. This increase is driven in part by the recognition that GRNs allow identification of biologically significant processes associated with a wide range of phenotypes that can be missed when looking at gene expression alone. Despite the utility of GRNs, published studies have generally failed to provide access to the GRNs themselves because the size of the inferred networks can exceed size limits for supplementary data allowed by journals and websites and because there have been no public repositories for these genome-scale models. Although readers of these studies could recreate the networks used in the analyses, the time and cost of inferring hundreds or thousands of large-scale networks at the sample level can be prohibitive. These difficulties with recreating the networks limit both assessment of the reproducibility of published studies and the use of the inferred GRNs for additional analyses.

GRAND represents a curated large-scale repository for genome-scale GRNs paired with extensive phenotypic information. In its current release, GRAND is populated with 12,468 GRNs and 173,013 targeting scores linking TFs and miRNAs to their target genes using a collection of GRN inference methods available in netzoo. These models were generated using data from large repositories including GTEx, TCGA, CCLE, and the Connectivity Map, as well as selected studies from GEO. The GRNs in GRAND are classified into four large groups—small molecule screens, cancer tissues, normal tissues, and cell lines. GRAND allows users to browse, visualize, analyze, and download these GRNs either through the web interface or programmatically through GRAND’s API. GRAND also allows network-based queries to identify small molecule candidate drugs that can potentially correct altered regulatory processes in disease states and users can upload their own networks to run the collection of tools in GRAND.

Future releases of GRAND will include additional gene regulatory network models from an increasing number of biological contexts, as well as networks inferred using newly developed inference methods designed to take advantage of the ever more complex multi-omics data that we can now generate. In addition, we will include models inferred from additional public data sets, including a larger number of cancer regulatory models and GRNs inferred from single-cell expression data. We also plan to include additional analytical tools and features requested by users of the resource.

## AVAILABILITY

GRAND is accessible at https://grand.networkmedicine.org and all source code is available at https://github.com/QuackenbushLab/grand.

## ACKNOWLEDGEMENT

The authors would like to acknowledge Tian Wang and Yunhao Huo for the assistance with graphical design, Dr. Alberto Noronha and the Quackenbush lab members for helpful discussions. Icons in figure 1 and figure 3 were used from Font Awesome using the following license https://fontawesome.com/license.

