## Supplemental file for "GRAND: A database of gene regulatory network models across human conditions"

### I. Supplementary figures

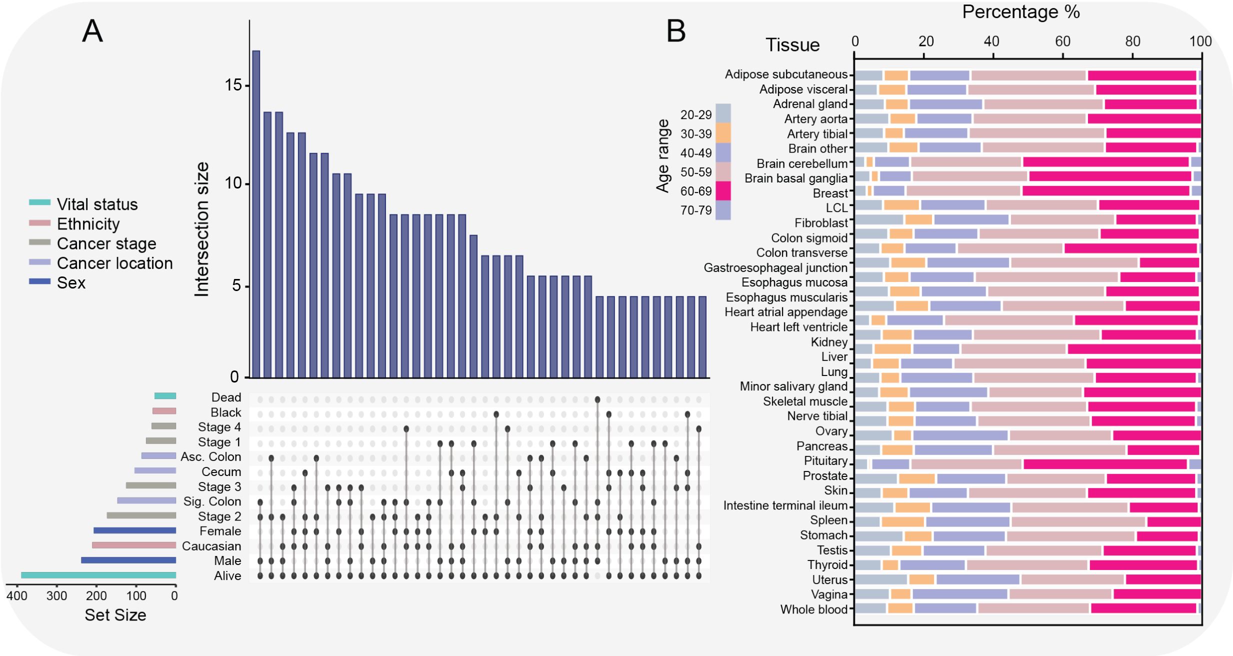


**Figure S1 - The small molecule resource integrates drug characteristics with cell line phenotypic information.** The resource combines information from the Connectivity Map and the Drug Repurposing Hub (DRH) for more than 173,013 samples. Each sample is represented by an edge in the diagram.

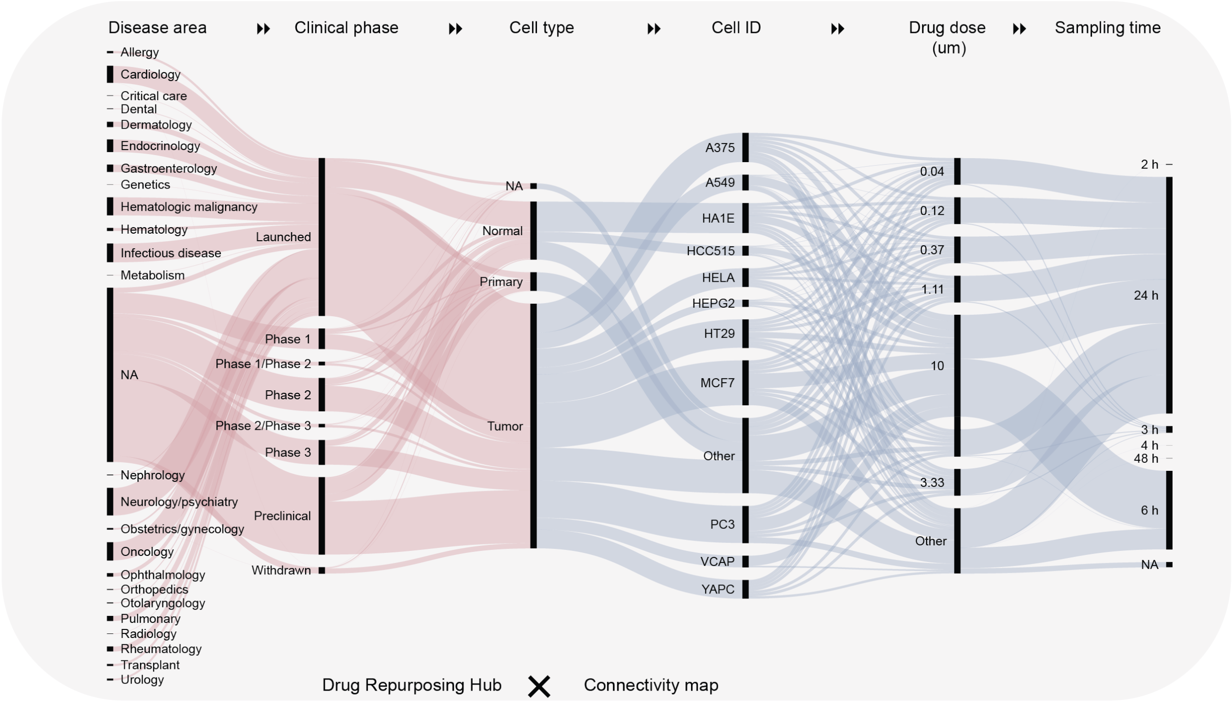


**Figure S2 - Summary statistics of the cancer resource and the tissues resource. A.** UpSet plot of the set intersection size of the most important clinical attributes in the cancer resource using colon cancer as an example. The plot represents the intersection between different groups of clinical attributes, for example the first group has 16 patients that belong to the groups “alive, “ “stage 2” cancer, and with the cancer located in the “sigmoid colon.“ **B.** Age distribution of the subjects from GTEx included in the tissues resource**.**

#### II. Supplementary tables

**Table S1 - Database content by condition and regulation modality. PAAD: Pancreatic adenocarcinoma, GBM: Glioblastoma multiforme, COAD: Colon adenocarcinoma.**

| Resource | Types | Data set | Regulation | Number of GRNs | Network type | Reference |
|---|---|---|---|---|---|---|
| **Cell lines** | LCL, Fibroblast | GTEx | TF | 2 | Aggregate | (1) |
| **Cell lines** | 35 tissues of origin | CCLE | TF | 1,376 | Single-sample | This paper |
| **Cell lines** | 35 tissues of origin | CCLE | miRNA | 1 | Aggregate | This paper |
| **Tissues** | 36 tissue types | GTEx | TF | 36 | Aggregate | (2) |
| **Tissues** | 36 tissue types | GTEx | miRNA | 36 | Aggregate | (3) |
| **Tissues** | 29 tissue types | GTEx | TF | 8,279 | Single-sample | (4) |
| **Cancer** | PAAD | TCGA | TF | 150 | Single-sample | (5) |
| **Cancer** | GBM | TCGA/GGN | TF | 1,023 | Single-sample | (6) |
| **Cancer** | COAD | TCGA/GEO | TF | 1,638 | Single-sample | (7) |
| **Cancer** | All | TCGA | TF | 22 | Aggregate | This paper |
| **Small molecules** | 2,858 labeled drugs | CLUE/DRH | TF | 173,013 | Targeting scores | This paper |
